# Activity in primate visual cortex is minimally driven by spontaneous movements

**DOI:** 10.1101/2022.09.08.507006

**Authors:** Bharath C. Talluri, Incheol Kang, Adam Lazere, Katrina R. Quinn, Nicholas Kaliss, Jacob L. Yates, Daniel A. Butts, Hendrikje Nienborg

## Abstract

Organisms process sensory information in the context of their own moving bodies, an idea referred to as embodiment. This idea is important for developmental neuroscience, and increasingly plays a role in robotics and systems neuroscience. The mechanisms that support such embodiment are unknown, but a manifestation could be the observation in mice of brain-wide neuromodulation, including in the primary visual cortex, driven by task-irrelevant spontaneous body movements. Here we tested this hypothesis in macaque monkeys, a primate model for human vision, by simultaneously recording visual cortex activity and facial and body movements. Activity in the visual cortex (V1, V2, V3/V3A) was associated with the animals’ own movements, but this modulation was largely explained by the impact of the movements on the retinal image. These results suggest that embodiment in primate vision may be realized by input provided by the eyes themselves.

---

Organisms process sensory information not in isolation but within the context of a moving body that is interacting with the environment, a phenomenon whose importance is underscored in developmental neuroscience^1^, and in robotics and artificial intelligence^2^, from vacuum-cleaning robots to self-driving cars^3^ (see also Fei-Fei, L, Montreal AI Debate 2: https://www.youtube.com/watch?v=XY1VTLRIsNo). A longstanding question in systems neuroscience is the degree to which this embodiment influences sensory processing^4–6^. In mice, locomotion affects neural activity in primary visual cortex (V1)^7–18^ and spontaneous movements are associated with pronounced brain-wide activity, including in V1^19–21^. The work in mice suggests that embodiment plays a crucial role in shaping processing in the visual cortex, although it is unclear whether similar phenomena are observed in other species^22–24^. The degree to which such movements influence responses in the primate visual cortex is of interest for several reasons. First, it could be a direct observation of embodiment that can be dissected into mechanisms and probed to understand its computational principles. Second, it addresses a fundamental question about the functional organization and degree of modularity of the primate cerebral cortex^23^. And third, it could have far-reaching implications for the interpretation of past neurophysiological studies of the primate visual system, in which the animals’ spontaneous body movements were not monitored.

Here, we ask whether the animal’s own body movements are associated with modulations of neural activity in visual cortex of macaque monkeys. We mirrored the experimental approaches used in studies in mice to facilitate the comparison between the data in mice and the data in primates: we used videography to monitor the animals’ movements^19,20^, and statistical modeling^20,21^ to relate the movements to neural spiking activity recorded in visual cortex (V1, V2, V3/V3A). Consistent with the results observed in mice, we found activity associated with the animals’ own spontaneous body movements. But when accounting for the fact that some of these movements also changed the retinal input to the neurons in visual cortex, this movement-related activity largely disappeared. As a model-free approach we also compared the modulation by spatial attention with the modulation by the animals’ own movements. The modulation by movement was an order of magnitude smaller than that by attention, and not associated with the modulation by attention. We conclude that in macaque early and mid-level visual cortex, activity is minimally driven by the animal’s own spontaneous body movements.

## Results

### The macaque monkeys move spontaneously while performing visual tasks

We used multichannel extracellular recordings targeting V1, V2 and V3/V3A combined with video-based monitoring of the body and face, in two alert macaque monkeys. The animals performed a visual fixation task or visual discrimination task (Fig. **1**). They fixated a spot on the center of the display during stimulus presentation epochs, which allowed us to reconstruct the stimulus in retinal coordinates. Outside of the stimulus-presentation epochs, the animals freely moved their eyes. Like the mice in the previous studies^19–21^, the monkeys were head-fixed, but otherwise free to move their arms, legs, and bodies throughout and in between stimulus presentations while seated. As the videography confirmed, the animals often fidgeted and moved spontaneously throughout the recording sessions (Fig. **1B**, Supplementary movie **S1**). To identify the animals’ movement patterns from the videos we used singular value decomposition (SVD), analogous to previous work in mice^19^ (Fig. **1B**). From these data, we could directly ask to what extent the animals’ own spontaneous face and body movements predict neural activity in the primate visual cortex.

**Fig. 1.**
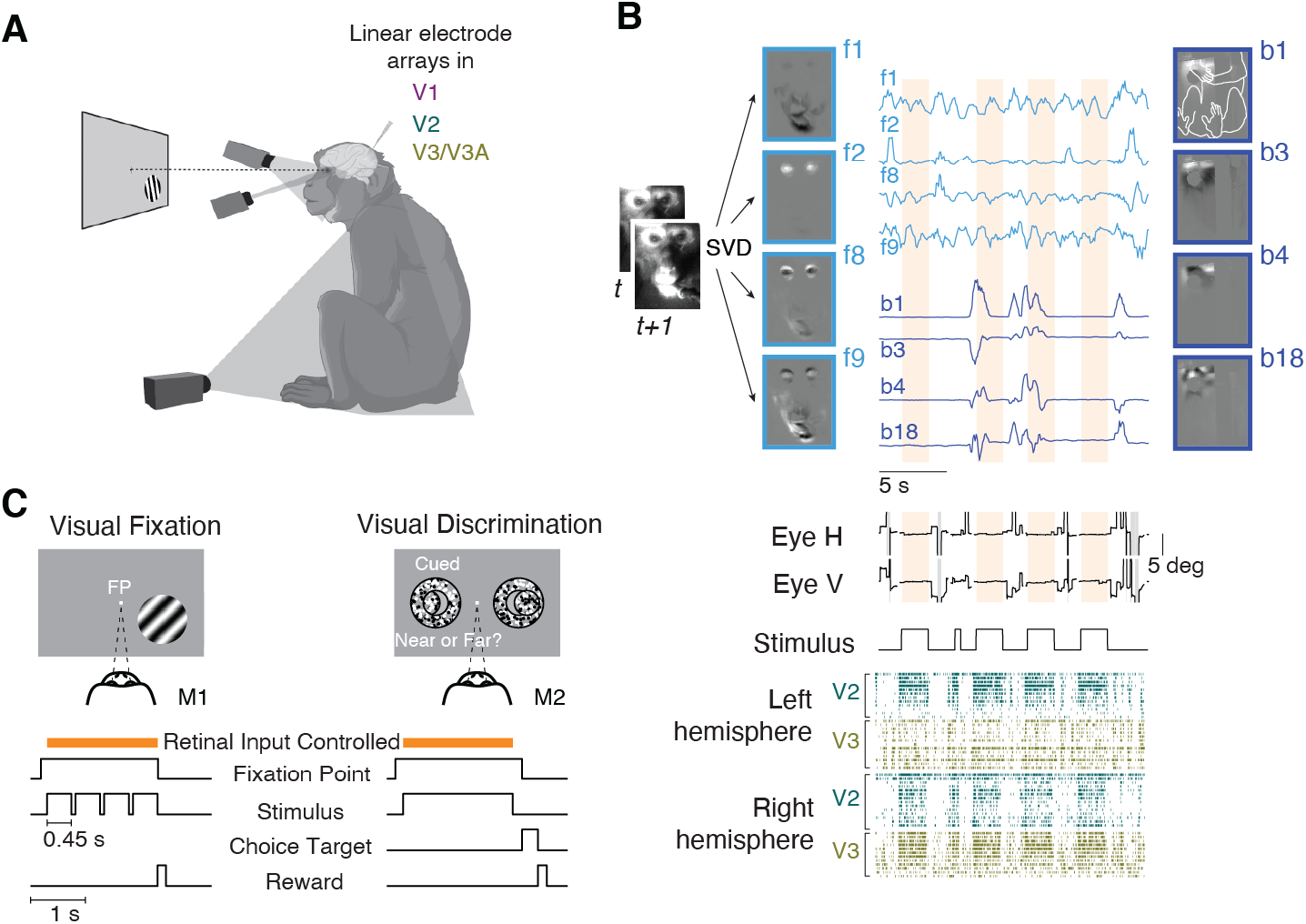
Monitoring spontaneous body movements during task performance in macaque monkeys. **(A)** The setup. The animals performed a visual task while extracellular activity in their visual cortex was recorded and the animals’ body, face, and eye movements were monitored via video, with one camera directed at the body, one at the face, and a video-based eye tracker. **(B)** Movements recorded by video (example from M2) were decomposed (singular value decomposition, SVD) generating multiple components of face and body movement that map onto, e.g., movements of the mouth (face component 1, f1), eye blinks (f2), combinations of face parts (f8, f9), and combinations of hand, arm, leg and body movement (body components b1, b3, b4, b18; outline of the monkey body shown in b1; grayscale shows normalized components; traces show normalized temporal profiles of the video projected onto the components); middle panels show eye positions and stimulus ON/OFF periods. Gray bands in eye position traces indicate interrupted eye signals due to blinks or eccentric eye positions. Bottom: Sample spike-rasters of simultaneously recorded units in the left and right hemisphere of V2 and V3/V3a. In each row spike times from one unit are shown as vertical ticks. **(C)** Animal M1 performed a visual fixation task. Animal M2 performed a visual discrimination task combined with block-wise manipulation of spatial attention. The retinal input was controlled during periods (orange bar) when the animals fixated on a fixation point (FP) at the center of the screen.

### Spontaneous movements predict neural activity when the retinal input is uncontrolled

We analyzed the data using a linear encoding model^20,21^, to predict the neural activity using a set of “predictors” (Fig. **2A**). The predictors include controlled variables in the experiment related to the task and the stimulus, and uncontrolled but observable variables such as the temporal profiles of the movement components (Fig. **2A**, labels on left of the panel), as well as temporally shifted versions of these predictors. The model successfully captures the stimulus-aligned response: the predicted firing rate at 16ms resolution and the peristimulus spike density function (SDF) over all trials are closely matched (Fig. **2A**, right, **2C**). Such peristimulus SDF-based validation, however, obscures the effects of spontaneous movements on both the model and the data, because the movements are not necessarily time-locked to times in the trial. Indeed, while some movements were aligned with the trial events, there was substantial movement variability throughout the trial, including the stimulus presentation period, when the animals maintained visual fixation on a small dot in the center of the screen (Fig. **2B**). Thus, to capture trial-to-trial variability that included the potential role of the animal’s own body movements we evaluated model performance on an individual trial level (percent variance explained: %VE, see Methods), for each of the 900 units across both animals and all areas (Fig. **2D**).

**Fig. 2.**
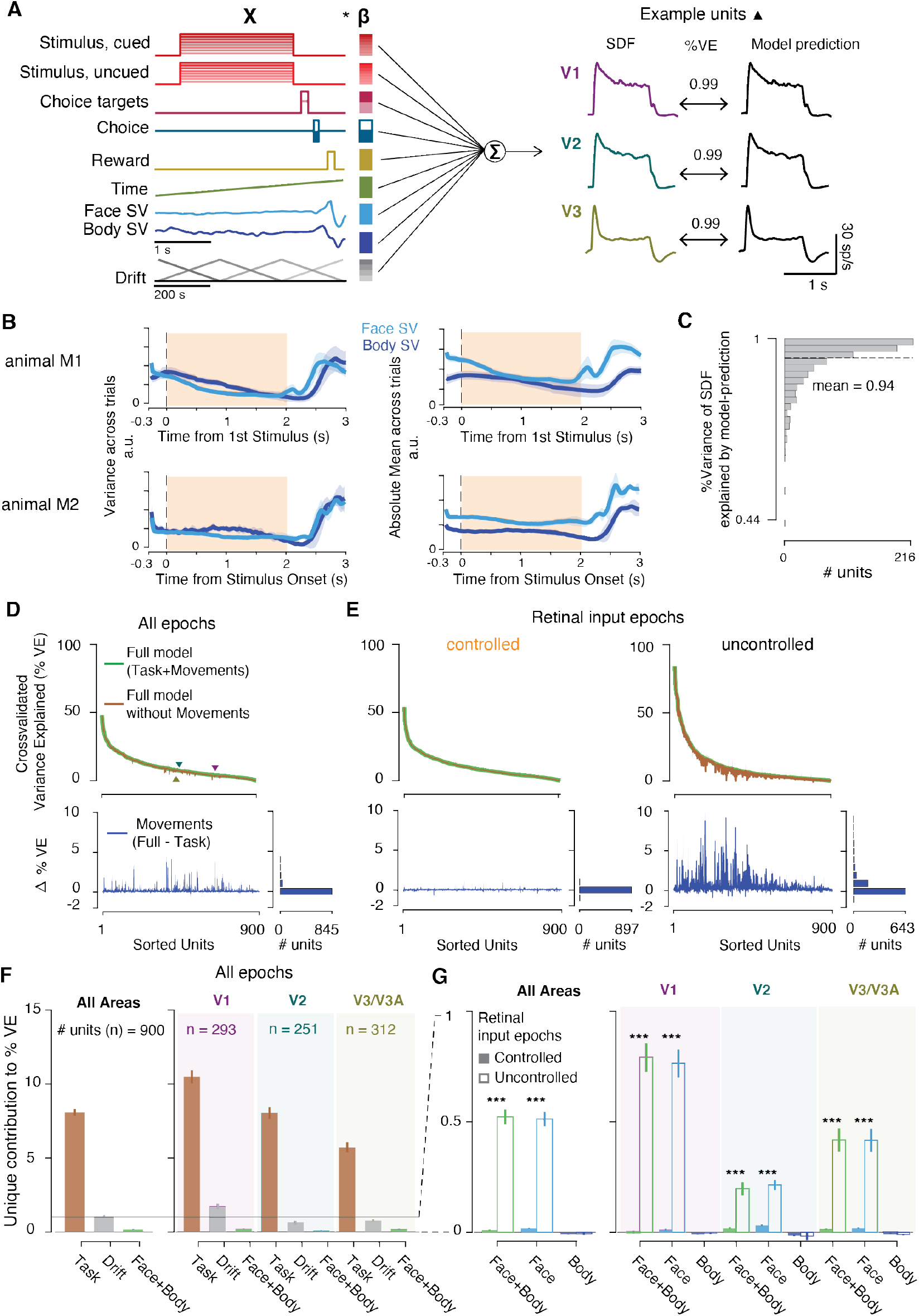
Body and face movement of the macaque monkey has minimal impact on neural activity in its visual cortex. **(A)** Linear encoding model predicts neural firing in visual cortex (the predictors, labels left, are for the task used in M2. For M1, see Supplementary Fig. **S2**. The three traces show the peristimulus SDFs for sample units in V1, V2, and V3/V3A (left) and the model predictions (right). **(B)** Mean variance (left) and absolute mean (right) of the top 30 face and body movement components across trials (M1, top; M2, bottom). Shaded error bars, standard deviation across sessions; shaded area: epochs during which the animals maintained visual fixation (controlled retinal input). **(C)** Histogram showing the distribution of cross validated variance explained (%VE, mean=94%, n=900 units from both animals) of the SDF by the model predictions across units. **(D)** Top: Variance explained across all timepoints by the model with (green), and without (brown) movement predictors for all units (%VE, mean = 9.8% and 9.67% respectively). Triangles show the example units from **(A)** (differences in %VE in **(A)** and **(D)** largely result from spike count variability at these high time resolutions). Bottom: Difference in variance explained by the two models, reflecting the %VE by movements. Units are ranked by their variance explained by the full model. **(E)** Same as **(D)**, but separately for epochs when retinal input was controlled (left, shaded interval in **B**), and not controlled (right). **(F)** Mean unique variance explained by different covariates towards the full model, for units across all areas (left; including 44 units for which the area could not be assigned) and separated by area (right). Error bars, standard error of mean across units. **(G)** Mean unique variance explained by movement covariates towards the full model, separately for controlled and uncontrolled retinal input epochs, for units across all areas (left), and separated by area (right). Note the smaller y-scale compared to **F**. Error bars, standard error of mean across units; ***, p < 0.001.

To address our central question of whether neural activity can be explained by the animal’s spontaneous movements, we compared two models: first, the full model, with all the predictors (Fig. **2D**, full model; green), and second, a “task-only” model (Fig. **2D**, brown), which was the full model but with the contribution of the movement predictors removed (see Methods). The difference in variance explained between these two models – which is equivalent to the “unique variance”^20^ of face/body movement components that we consider below – is a measure of the amount of variance that can be explained uniquely by knowing the animals’ own movements (Fig. **2D**, bottom). The results show that activity in the primate visual cortex was predictable from the animal’s own movements, although the size of this contribution in the macaques was smaller compared with that seen in mouse visual cortex^20,21^.

To better understand how the monkeys’ own movements could impact neural activity in the visual cortex, we examined the amount of unique variance during different epochs of the trial: when the retinal input was controlled because the animal maintained visual fixation (orange bar, Fig. **1C**), and when the retinal input was uncontrolled. In the first type of epoch, the retinal image (gray screen or the stimulus) is known, and the corresponding predictors can contribute systematically to the model predictions. In the second type of epoch the retinal image is not known and thus could drive activity in a way that is predicted by movements causing these changes in retinal input. For each unit, we applied a threshold to determine if the neural activity of the unit was associated with face or body movement (threshold: unique variance >0.1%VE). Despite the fact that the animals moved spontaneously during both kinds of epochs (Fig. **2B**), the contribution of the model attributed to the movement almost completely disappeared when the retinal image could be inferred (Fig. **2E** left, 5% of units crossed the threshold; V1: 15/293, V2: 16/251, V3/V3A: 13/312), compared to when the retinal input was uncontrolled (Fig. **2E** right, 67% of units crossed the threshold, V1: 246/293, V2: 131/251, V3/V3A: 191/312). This result was robust for different thresholds of unique variance (table **S1**), and the increase in unique variance explained by movements when the retinal input was uncontrolled was significant (p<0.001 for each area and combined across areas, permutation test; Fig. **2G**; similar in each animal individually, Fig. **S3**). These results suggest that the larger unique contribution of the animal’s own movements when the retinal image was uncontrolled was the result of changes in the retinal image associated with these movements.

### Retinal input control reduces activity predicted by spontaneous movements

To validate this explanation, we compared the unique variance explained by movements inferred from the face view versus the body view. The explanation predicts that face movements, such as blinks or eye movements, are more likely than body movements to modulate neural activity. Consistent with this prediction, the increase in unique variance during epochs when the retinal input was uncontrolled was only significant for movements of the face (p<0.001 for each area and combined across areas, permutation test; Fig. **2G**). Moreover, removing the region of the eye from the face view reduced the increase in unique variance for retinal-input-uncontrolled epochs (p < 0.001 for each area and combined across areas, permutation test). Conversely, the contribution by body movements was small throughout all epochs (Fig. **2G**, unique variance due to body covariates: mean across epochs and units =-0.005 %VE, p=0.07), mirroring previous findings in mice^20^.

The data presented here suggest that accounting for retinal input removes the variability of neural responses that was predictable from the monkey’s own movements. To further test this idea, we analyzed the time-points during which the retinal input was uncontrolled, i.e., when the animals could move their eyes freely. We classified these time-points into two subsets. The first subset are times when the retinal input to the receptive fields of the recorded neurons could be inferred from the eye position, i.e., when the receptive fields were on a blank gray screen. The second subset are times when the retinal input to the receptive fields could not be inferred, e.g., when the gaze of the animal took receptive fields off the screen, and they likely included visual structure from the room. If the absence of retinal image control can explain the apparent neural modulation by body/face components, then the neural modulation by the animal’s movements should be higher in the latter case: when the retinal image is not known. This is exactly what we found (supplementary Fig. **S4**). Together, these results support a relationship between spontaneous movements in primates and visual cortical activity because of their correlation with changes in the retinal input.

### Attentional modulation is not associated with modulation by movements

Modulation by locomotion in mice shows parallels to the modulation by spatial attention in primates^25,26^. To therefore test for a potential relationship between neural modulation by spontaneous movements and by attention, we trained one animal to perform a visual discrimination task while manipulating spatial attention and monitoring the animal’s own movements. We observed the characteristic^27^ increase in neural response when the animal’s attention was directed to the receptive fields, including modest modulation by spatial attention in V1^28^ (Fig. **3A**), but this attentional modulation was not correlated with spontaneous body movements (Fig. **3B**, p>0.3 for all areas; neural modulation by spatial attention was also not correlated with the absolute value of the neural modulation by face/body movements, p>0.2 for all areas). The analysis in Fig. **3B** is model free, and shows that modulation by the animal’s own movement was about an order of magnitude smaller than the modulation by spatial attention (mean±standard deviation *MI*= 0.007±0.016, 0.009±0.03, 0.009±0.04; *AI* = 0.05±0.05, 0.11±0.07, 0.11±0.09 for V1, V2, V3/V3A, respectively; the distributions for *MI* versus *AI* differed significantly in all areas, p=0.003, p=10^−36^, p=10^−24^, for V1, V2, V3/V3A, respectively, t-tests, corrected for multiple comparisons). These findings corroborate our model-based results and provide evidence against an association in macaques between a modulation by an animal’s own movements and the modulation by spatial attention.

**Fig. 3.**
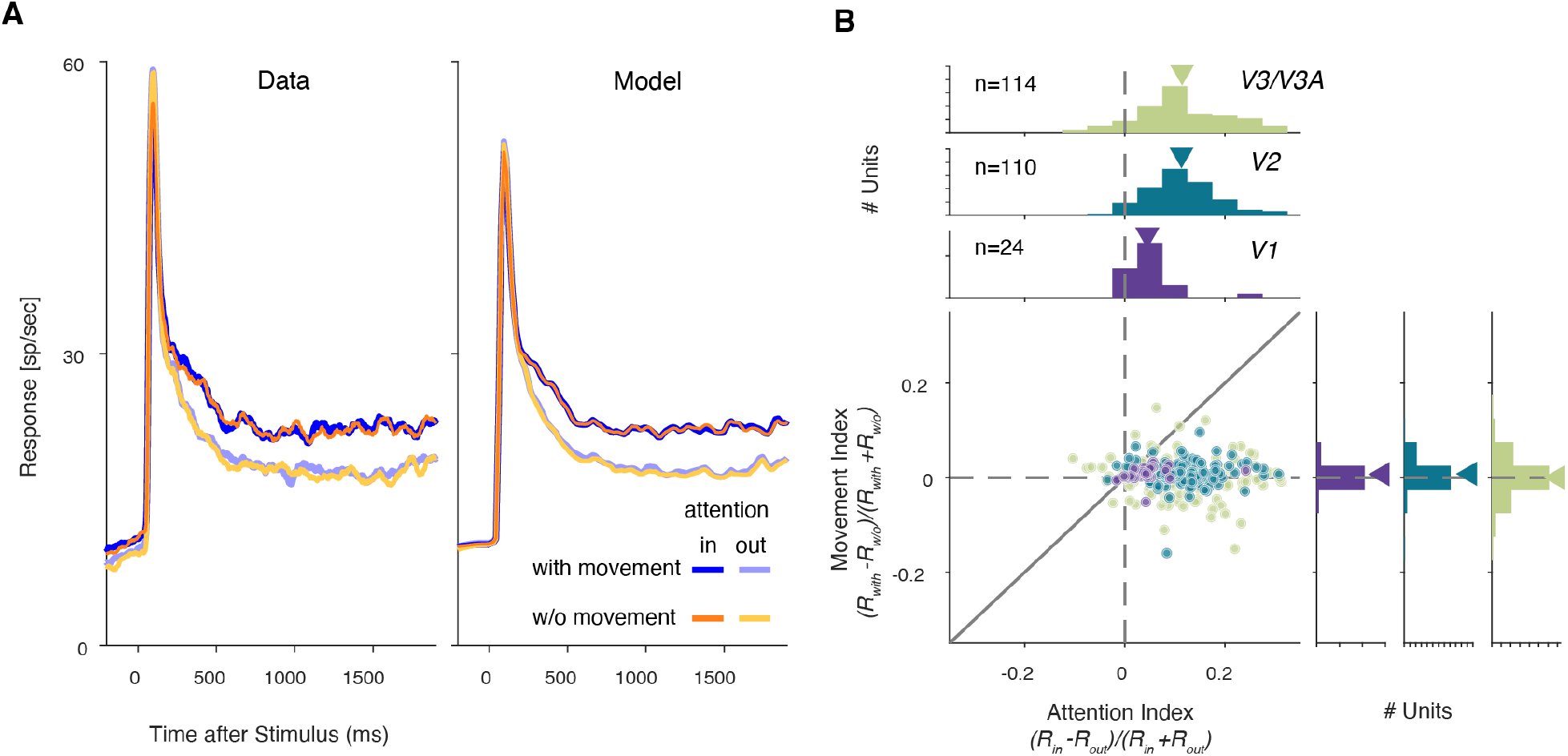
Modulation by spatial attention is not associated with modulation by movement. (**A**) Average stimulus-driven responses across all units (n=248, left; peristimulus SDF; right: rates predicted by the full model) separated by attention and the presence or absence of the animals’ spontaneous movements. (**B**) Modulation indices for attention (*AI)* are not correlated with those for movement (*MI*) in V1, V2 and V3/V3A.

## Discussion

The present results show that in macaque monkeys, spontaneous body and face movements accounted for very little of the variability of single-trial neural dynamics in macaque V1, V2 and V3/V3A. This contrasts with results in mice, where substantial modulation of visual cortical activity is associated with the animal’s own spontaneous movements^19–21^. The difference in results across species cannot be attributed to a difference in analysis methods: the present analysis was designed to replicate the approach used in mice (see Methods), and, when retinal input in the macaque monkeys was uncontrolled, spontaneous movements did account for appreciable neural variability, as in mice. Moreover, the neural measurements used presently recovered the expected modest levels of neural modulation caused by spatial attention^27,28^, even in V1, implying sufficient sensitivity of the neural recordings, and the present results parallel recent observations in marmosets of a quantitative difference in the neural modulation in visual cortex with locomotion between primates and rodents^29^.

Our results are also consistent with previously observed modulations by eye-movements, including microsaccades^30–37^, or gaze position^38^ but reveal that these are small (Fig. **2**, Fig. **S4**) compared to the overall response variability in the macaque visual cortex, in line with early reports^39,40^. The results here, combined with the findings in marmosets^29^, therefore suggest that decades of neurophysiological research on the primate visual system need not be revisited in light of the pronounced modulations by spontaneous movements observed in mice.

While the results here raise the possibility that some fraction of the neural modulation associated with movement observed in mice is related to uncontrolled retinal input, there are good reasons to suspect genuine differences in the mechanisms of embodiment between mice and monkeys. Primates and rodents differ not only in body anatomy, but also in behavior and brain organization. Primary visual cortex in mouse receives substantial direct projections from premotor areas^41^ but does not in monkey^42^, and the neuromodulatory system in visual cortex also differs in the two species^43^. A direct modulation of visual cortical responses by movement may be evident in higher visual areas in primates, which are perhaps a closer analogue of primary visual cortex in rodents^44–46^. The difference in results between mice and primates may therefore reflect corresponding differences in anatomy and behavior^47^. Primates must make sense of the statistics of their visual input and how that input is shaped not only by their body’s own locomotion^48^ but also prominently by their eye movements^49–51^. These demands may have selected mechanisms to emphasize embodiment that exploit input provided by the eyes themselves.

## Acknowledgments

We would like to thank the members of the laboratory of sensorimotor research for their insightful comments, Richard Krauzlis and Bevil Conway for discussions and comments on an earlier version of this manuscript and Florian Dehmelt, Paria Pourriahi and Lenka Seillier for early technical support.

## Funding

National Eye Institute Intramural Research Program at the National Institutes of Health 1ZIAEY000570-01 (HN)

German Research Foundation (DFG), 276693517 (TP6) (HN)

National Science Foundation, IIS1350990 (DAB)

## Author contributions

Conceptualization: JY, HN

Methodology: BCT, IK, JY, DAB, HN

Investigation: IK, KRQ

Data analysis: BCT, IK, AL, JY, HN

Video labeling: NK

Visualization: BCT, IK, HN

Funding acquisition: HN

Project administration: HN

Supervision: JY, DAB, HN

Writing – original draft: BCT, IK, HN

Writing – review & editing: BCT, IK, JY, DAB, HN

## Competing interests

Authors declare that they have no competing interests.

## Supplementary Information

### Materials and Methods

#### Animals

Two adult male rhesus monkeys (*Macaca mulatta*) were used as subjects (animal 1 (M1); animal 2 (M2); each 9 kg). All protocols were approved by the National Eye Institute Animal Care and Use Committee (M1) or by the relevant local authority (M2), the Regierungspräsidium Tübingen, Germany, and all experimental procedures were performed in compliance with the US Public Health Service Policy on humane care and use of laboratory animals. Under general anesthesia, the monkeys were surgically implanted with a titanium head post, and in a subsequent procedure with a recording chamber (19 mm inner diameter, cilux, Crist Instrument, Hagerstown, MD) over right hemispheric V1 (M1), and with two titanium recording chambers (25mm inner diameter) over the operculum of V1 on both hemispheres (M2), guided by structural MRI scans of the brain.

#### Behavioral tasks

##### Visual Fixation

Animal M1 was required to fixate on a small spot (Fixation Point (FP), 0.3° diameter) at the center of the screen for about 2sec to receive a liquid reward, while a drifting sinusoidal luminance grating was flashed four times (450 ms duration each separated by an interval of approximately 50ms of a blank screen) over the receptive fields (RFs) of the recorded units (left panel of Fig. **1B**). In addition to visual fixation, animal M2 also performed a visual discrimination task.

##### Disparity Discriminatio

Animal M2 performed a disparity discrimination task (right panel of Fig. 1B) previously described in detail^52^. Briefly, once the animal fixated on a FP (0.1° diameter), two circular dynamic random-dot stereograms (RDSs, for details see *Visual Stimuli*), consisting of a disparity-varying center surrounded by an annulus fixed at zero disparity, were presented, one in each visual hemifield. Stimuli presented in one hemifield were task-relevant. The animal had to judge whether the center disparity of the relevant RDS was protruding (‘near’; negative disparity) or receding (‘far’; positive disparity) relative to a surrounding annulus. After two seconds, the FP and the RDSs were replaced with two choice icons (circular RDSs at 100% disparity signal, one at the near, the other at the far signal disparity) positioned above and below the FP but horizontally offset towards the cued side. The animal was rewarded after making a saccade within 2sec after the onset of the choice icons, to the choice icon that had the same sign of the disparity signal as the stimulus. The task-relevant hemifield was cued by three instruction trials at the beginning of each 50-trial block. On instruction trials a single stimulus was presented on the task-relevant side. The vertical position (∼3° above or below the fixation point) of the choice icons was randomized across trials to prevent a fixed mapping between the chosen disparity sign and saccade direction.

#### Visual Stimuli

Visual stimuli were back-projected on a screen (Stewart Filmscreen, Torrance, CA) by a DLP LED projector (Propixx, VPixx Technologies, Saint-Bruno, QC, Canada; 1920×1080-pixel resolution). The display was achromatic, and the luminance steps were linearized (mean luminance: 72 cd/m^2^ for M1, 30 cd/m^2^ for M2). Visual stimuli were presented on a uniform display at the mean luminance. Separate images were delivered to the two eyes (120 Hz for M1 and 60 Hz for M2, for each eye) using a combination of an active circular polarizer (DepthQ, Lightspeed Design Inc., Bellevue, WA) in front of the projector and two passive circular polarizers with opposite polarities (American Polarizers, Reading, PA) in front of the eyes. The viewing distance was 45 cm for M1 and 97.5 cm for M2, at which the display subtended 74° by 42° for M1 and 32° by 18° for M2.

Stimuli used in the fixation task for M1 were drifting circular sinusoidal luminance gratings whose position and size were tailored to the collective RFs of the recording site. The spatial frequency was adjusted inversely proportional to the RF size and the temporal frequency was typically 4 or 5 Hz. The contrast of the stimulus during each of four 450-ms stimulus epochs on a trial was randomly chosen from 4 values (0, i.e., blank stimulus, 6.25, 25, and 100%) with equal probabilities.

Stimuli used in the disparity discrimination task for M2 were circular dynamic RDSs (50% black, 50% white dots, dot size typically 0.08° radius, 50% dot density) with a disparity varying central disk (3-5° in diameter, approximately matching the RF size of the recorded units) surrounded by an annulus of zero disparity (1° width). The positions of the dots were updated on each frame. The central disk consisted of signal frames randomly interleaved with noise frames. For each session, the signal disparities (one near disparity, one far disparity) were fixed. The center disparity of the stimulus was updated on each video-frame. On “signal frames”, the center disparity was one of the signal disparities, held constant across each trial. On a “noise frame”, the disparity of the center disk was randomly chosen from a uniform distribution of 9 values equally spaced from -0.4° to 0.4°. The task difficulty on a trial was defined as the ratio of the signal to noise frames such that 100% means that all frames were signal frames, and 0% means all frames were drawn from the noise distribution. On a 0% trial, the reward was randomly given 50% of times. The choice target icons were also circular RDSs but slightly smaller than the stimuli, and always presented at 100% near and far signal. We assessed disparity tuning before the behavioral task in separate visual fixation experiments using RDSs (450 ms duration), whose disparity varied typically from -1° to 1° in 0.1° increments. The two signal disparities in each session were chosen to approximately match the preferred and non-preferred disparities by most of the recorded units.

Receptive fields (Supplementary Fig. **S1**) of the recorded units were first approximated by a bar stimulus whose orientation and position were manually controlled, then quantitatively measured with strips of horizontal or vertical bars (450 ms duration each, typically white and black bars but sometimes RDSs at the preferred disparity when they evoked stronger responses) that were equally spaced over the range covering RFs estimated by manual sweeping (typically 9 to 11 positions whose intervals were determined by the collective RF range).

Visual stimuli were generated in MATLAB (MathWorks, Natick, MA) by custom-written code^53^, adapted from Eastman & Huk^54^ using the Psychophysics toolbox^55^.

#### Electrophysiological Recordings

Extracellular recordings were made from areas V1, V2 and V3/V3A using multi-channel laminar probes (Plexon Inc., Dallas, TX; V/S Probes, 24/32 channels, 50 to 100 μm inter-contact spacing). Neuronal signals were amplified, filtered (250 Hz to 5 kHz) and digitized (30 kHz sampling rate) by the Grapevine Neural Interface Processor (NIP, Ripple Neuro, Salt Lake City, UT) run by the Trellis software (Ripple Neuro, Salt Lake City, UT) that interfaced with MATLAB via Xippmex (v1.2.1; Ripple Neuro, Salt Lake City, UT).

We inserted recording probes on each day of experiments via the operculum of V1 using a custom-made (M1) or customized (M2; NaN Instruments, Israel) micro-drive placed approximately normal to the surface. We initially mapped the recording sites using single tungsten-in-glass electrodes (Alpha Omega, Nazareth, Israel) to determine the receptive field locations and assess the selectivity for horizontal disparity. During data collection, visual areas were identified using two physiological criteria: 1) transitions from a gray to white matter, which was typically characterized by a silent zone that spanned a few consecutive channels showing weak or no visually driven responses, 2) abrupt shifts in the receptive field location and size, and often abrupt changes in the tuning preferences for orientation or disparity. Final assignments of channels to visual areas were done offline with the aid of receptive field maps constructed from receptive field location and size determined from quantitative fitting (see below) across all sessions (see supplementary Fig. **S1**), combined with the structural MRI scans. Because of the similarity between the disparity selectivity in V3 and V3A^56^, we did not seek to further assign channels to V3 or V3A, and instead designate them collectively as V3/V3A.

On each day of experiments, after the laminar probe was advanced to a depth at which most channels spanned the visual area we intended to record from, we usually advanced it further down to confirm the visual area underneath. Then, we withdrew the probe back to the desired depth and waited for at least 30 min before data collection to allow time for the tissue around the probe to be stabilized, thereby to minimize vertical drifts of the recording site along the probe. We mapped the receptive fields before, sometimes in between, and after data collection to diagnose drifts of the neural tissue relative to the electrode via the receptive field position across the channels during data collection. We only included units that remained in the same visual areas during the entire data collection period and excluded units whose activity was picked up by channels positioned within the transition depth between visual areas at any time during data collection.

#### Measurements of eye position

We monitored the animals’ binocular eye positions using the EyeLink 1000 infrared video tracking system (SR Research, Ottawa, ON, Canada) at a sampling frequency of 500 Hz.

#### Recording of face and body movements

To record the face and body movements of the animals during data collection, we installed infrared (940nm) LEDs and at least two cameras (Fig. **1A**; M1 – Stingray camera integrated in a CinePlex Behavioral Research System, Plexon Inc., Dallas, TX, 60 or 80 Hz sampling rate, downsampled to 20Hz and spatially downsampled by 2×2 pixels for analysis; M2 – Imaging Source DMK camera; triggered image acquisition at 12.5 Hz), one pointing to the face, and one to the front view of the body.

### Data Analysis

#### Spike sorting

We sorted spikes from single- or multi-units offline using Kilosort2.5^57^ followed by manual curation in Python (www.github.com/cortex-lab/phy) for data from M1, and using the Plexon Offline Sorter (v3.3.5; Plexon Inc., Dallas, TX) for data from M2. We analyzed spikes from both single- and multi-units isolated by the spike sorting procedures, which we refer to as units without distinction.

#### Receptive fields

To measure receptive fields, we averaged the multi-unit response (spike count during stimulus interval) on each recording channel for each position of the bar stimuli. We fit a Gabor function to the mean response as a function of stimulus position, separately for the horizontal and vertical dimensions, using MATLAB (*lsqnonlin*). The center of the receptive field was defined as the position at the peak of the fitted function, and the width as the distance between the two positions flanking the peak at which the fitted function reached 20% of its peak above the offset (Fig. **S1**).

#### Motion decomposition

To quantify the face and body movements, we selected regions of interest (ROI) from the videos with the face view and the frontal body view to include only the animal’s face and body. The movements in the selected ROIs were decomposed into movement components using singular value decomposition (SVD) following the method in Stringer at al.^19^ (www.github.com/MouseLand/FaceMap), via temporal-segment wise SDV (∼1 min long segments of the videos) (Fig. **1C**). The motion matrix *M* of the video, where *M* is the absolute pixel-wise difference between two consecutive frames (number of the pixels in the ROI × number of the video frames minus 1), was then projected onto the first 1000 movement components to calculate their temporal profiles. These temporal profiles correspond to the face/body movement regressors used in the ridge regression modeling approach described below. To evaluate the contribution of the movement components of the eye region in the face view to neural modulation (see Results), we performed the same SVD analysis on the face videos after the eye regions were removed from the face ROI.

#### Modeling neural activity during trials

We modeled the spiking activity of each unit as a linear combination of task-related and task-unrelated events within a session using ridge regression adapted after Musall, Kaufmann et al.^20^. Our linear multivariate regression is thus analogous to the approach used previously in mice^20,21^. Although a non-linear model might achieve better overall predictions, we used a linear statistical model to facilitate this comparison to mice, such that it cannot account for the discrepancy between the previous findings in mice and our findings in the macaque visual cortex.

Regressors for task-related events reflect the stimulus, the time since the beginning of the trial, the timing of reward in both animals, and additionally, the presence of choice-targets and saccadic choice in animal M2 (Fig. **2A**). Regressors for task-unrelated events were based on face and body movements, and a slow drift term to capture non-stationarities in firing rates of each unit. Below we describe the individual regressors:

##### Stimulus regressors

stimulus regressors were discrete binary vectors with one dimension for each distinct stimulus (i.e., disparity and contrast). They had the value 1 for the appropriate stimulus dimension in the time periods spanning the stimulus presentation window and 0 elsewhere. Separate regressors were used to model different stimulus values. In addition, in animal M1, within each stimulus value (i.e., the four contrast values), separate regressors were used for the 4 successive samples in time (supplementary fig. **S2**). In animal M2, within each stimulus value (i.e., the disparity value on each video-frame), separate regressors were used for stimulus presentations on the left and right hemifields (i.e., whether the attended stimulus was within or outside of the receptive field of the recorded neuron). This allowed us to capture modulation of spiking activity as a function of sample position within the stimulus sequence and stimulus contrast in animal M1, and as a function of disparity and attended location in animal M2.

##### Reward regressors

reward regressors were discrete binary vectors with value 1 at reward onset and 0 elsewhere.

##### Time regressors

time regressors were discrete binary vectors with value 1 at stimulus onset and 0 elsewhere and were used to model modulations in spiking activity due to stimulus onset and offsets (see fitting procedure).

##### Choice-target, and Choice-saccade regressors

animal M2 performed a discrimination task requiring him to make a saccade to one of the two targets presented after the stimulus offset. Target regressors were discrete binary vectors with value 1 at target presentation and 0 elsewhere. Separate regressors were used to model targets presented offset to the left and right hemifields. Choice regressors were discrete regressors with the value +/-1 to model saccades to the top and bottom target when the animal reported the choice and 0 elsewhere.

##### Drift regressors

non-stationarity in firing rates for each unit was modeled as a set of analog regressors using tent basis functions spanning the entire session^52^. These basis functions allow for a smoothly varying drift term that can be fitted as linear model terms. We defined anchor points placed at regular intervals within each session (10 and 8 anchor points for animals M1 and M2, respectively), each denoting the center of each basis function. The basis function has a value 1 at the corresponding anchor point, and linearly decreases to 0 at the next, and previous anchor point, and remains 0 elsewhere. Thus, any offset at each timepoint due to slow drift in firing rate is modeled by a linear combination of the two basis functions. While the drift regressors were included to account for non-stationarities related to experimental factors, they would also capture factors related to slowly changing cognitive states throughout a session^58^. To therefore avoid that the drift predictors accounted for the block-wise alternation in spatial attention for M2, we ensured that no more than one anchor point was used for each pair of successive, i.e., alternating, blocks of attention.

##### Face & Body movement regressors

the temporal profiles of the top 30 SVD components (SVs) of videos capturing movements in the face and body regions in both animals were used as analog regressors to model modulation in spiking activity due to movements. Note that since we did not additionally include regressors for pupil-size or eye-position, this gives the included movement regressors the possibility to also explain neuronal variability that might otherwise be explained by pupil regressors due to the correlation between these covariates^16^. To avoid overfitting, we limited our analysis to 30SVs, but our results were qualitatively similar when the top 200 SVs were used instead (supplementary fig. **S5**).

#### Fitting procedure

Recordings from each session were first split into individual trials. We modeled only successfully completed trials. Each trial was defined by a 300ms pre-stimulus period, the stimulus presentation window, and a 1000ms window after stimulus offset. This allowed us to split time-periods within an individual trial into those where the retinal input was controlled, i.e., the animal maintained visual fixation, and where the retinal input was uncontrolled. Time points within each session were discretized into non-overlapping 16.67ms wide time bins, matching the lower framerate of the stimulus displays used for the two monkeys. Spiking activity of each unit was quantified as the number of spikes in each time-bin, and all the regressors were down-sampled to 60 Hz while preserving their discrete/analog nature. On trials where the 1s post-stimulus window of the current trial overlapped with the 0.3s pre-stimulus window of the next trial, we reduced the post-stimulus window to only include the non-overlapping time bins.

Because the effect on neural activity of a given regressor will often play out across time, we modeled the effect of each regressor using a time-varying “event kernel” by creating numerous copies of individual regressors each shifted in time by one frame^20^ relative to the original using pre-defined time windows. These time-windows for stimulus, reward, and choice-target regressors were 250ms post-event, for choice-saccade regressors were 500ms pre- and post-event, and for time regressors spanned the entire duration of the trial following the stimulus onset including the post-stimulus window. The time-varying kernels of the analog movement regressors were modeled by convolving the temporal profiles of the corresponding component with separate tent basis functions with anchor points at -100ms, 0ms, and 100ms with respect to the movement event. This allowed us to capture the temporal dependence of spiking activity on the movement within a 400ms time window, resulting in a total of 90 regressors each for face and body movement components. All the event kernels were constructed at the level of individual trials.

We fit the models using ridge regression and 10-fold cross-validation across trials to avoid overfitting. Trials were randomly assigned to training or test dataset within each fold such that no event kernel spanned samples from both the training and test sets. Separate ridge penalty parameters were estimated for each unit during the first cross-validation fold which were then used in subsequent folds.

#### Model performance

We used cross-validated variance explained (%VE) as the measure of model performance. This is computed based on the variance of the residual of the model prediction (prediction minus the binned spike count) compared with the overall variance of the observed binned data. Note that %VE at the single-trial level at these time resolutions (16ms bins) is dominated by spike-count variability, and the same models that explained on average 94% of the variance in the SDF averaged across trials (Fig. **2C)** explain a mean of 9.8% VE (Fig. **2D**). Furthermore, to determine the “unique” effect of different task-related and task-unrelated events on the spiking activity, we estimated the “unique variance” as defined by Musall, Kaufmann et al.^20^. This metric was devised to account for the fact that many predictors in the model are correlated. It is the variance explained by each class of regressors by computing the %VE for a reduced model obtained by shuffling in time only the regressors under consideration leaving all the others intact and subtracting this from the %VE of the full model. Note that by shuffling rather than eliminating a given regressor, the resulting model will have the same amount of parameters as the full model and thus, if the regressor contained no additional (or “unique”) information to predict the neural response, it would result in the same %VE, The resulting difference (Δ %VE) thus gives a measure of the predictive power unique to each regressor^20^.

#### Movement and Attention Index

To determine periods with movement (Fig. **3**) we used the motion matrix *M (*see section *Motion decomposition)* for the face and body, where *M* is the absolute pixel-wise difference between two consecutive video frames (number of the pixels in the ROI × number of the video frames minus 1). We then averaged *M* over pixels to compute the average motion versus time 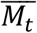 (1x number of frames minus 1). Periods with movement were defined as those when 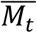 exceeded the 80th percentile across all time-points of 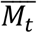 in either the face or body view, while periods without movement were defined as those for which 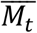 was below its median across all time-points, in either the face or body view. (We note that we confirmed that the results were qualitatively similar when we used 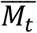 from only the body view or only the face view, indicating that neither type of movement had a sizable effect on *MI*.) We calculated the movement index (MI) and attention index (AI) based on the average spike rates (*R;* computed after removing non-stationarities across the recording session using the drift-term of the linear regression model described below) from 0.15-2sec after stimulus onset, as 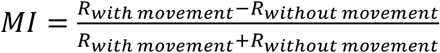 and 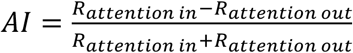.

We computed the spike density functions (Fig. **3A**) by convolving peri-stimulus time histograms (1ms resolution) for each unit with a temporal smoothing function (half Gaussian function; standard deviation 30ms) and averaging this across units.

#### Dataset

Our dataset consists of a total of 1407 units: 1139 units from M1 recorded in 54 sessions and 268 units from M2 recorded in 5 sessions. We excluded 507 units from the analysis that failed to meet the following criteria: (1) a minimum mean firing of 2 spks/s during stimulus presentations epochs in each of the four quartiles of the session, and (2) a minimum of 0 %VE of the full model during both retinal input controlled and uncontrolled epochs. Among the remaining 900 units, 653 units were from M1 (V1 - 269, V2 - 143, V3/V3A - 198) and 247 units were from M2 (V1 - 24, V2 - 108, V3/V3A - 114). Results were qualitatively similar when the minimum firing rate criterion was relaxed to include 1343 units in the model (Supplementary Fig. **S6**). For the model-free analysis in Fig. **3**, we only used the first criterion, avoiding sub-selection of units based on model-fits. We did not assign visual areas to 44 units recorded in three sessions from M1 in which the receptive location and size were not consistent with the overall topography of the offline receptive field map as to unambiguously assign the recording sites but included them when data were combined across areas.

#### Statistical tests

We used nonparametric permutation tests^59^ to test for group-level significance of individual measures, unless otherwise specified. This was done by randomly switching the condition labels of individual observations between the two paired sets of values in each permutation. After repeating this procedure 10,000 times, we computed the difference between the two group means on each permutation and obtained the P value as the fraction of permutations that exceeded the observed difference between the means. All P values reported were computed using two-sided tests unless otherwise specified.

**Fig. S1.**
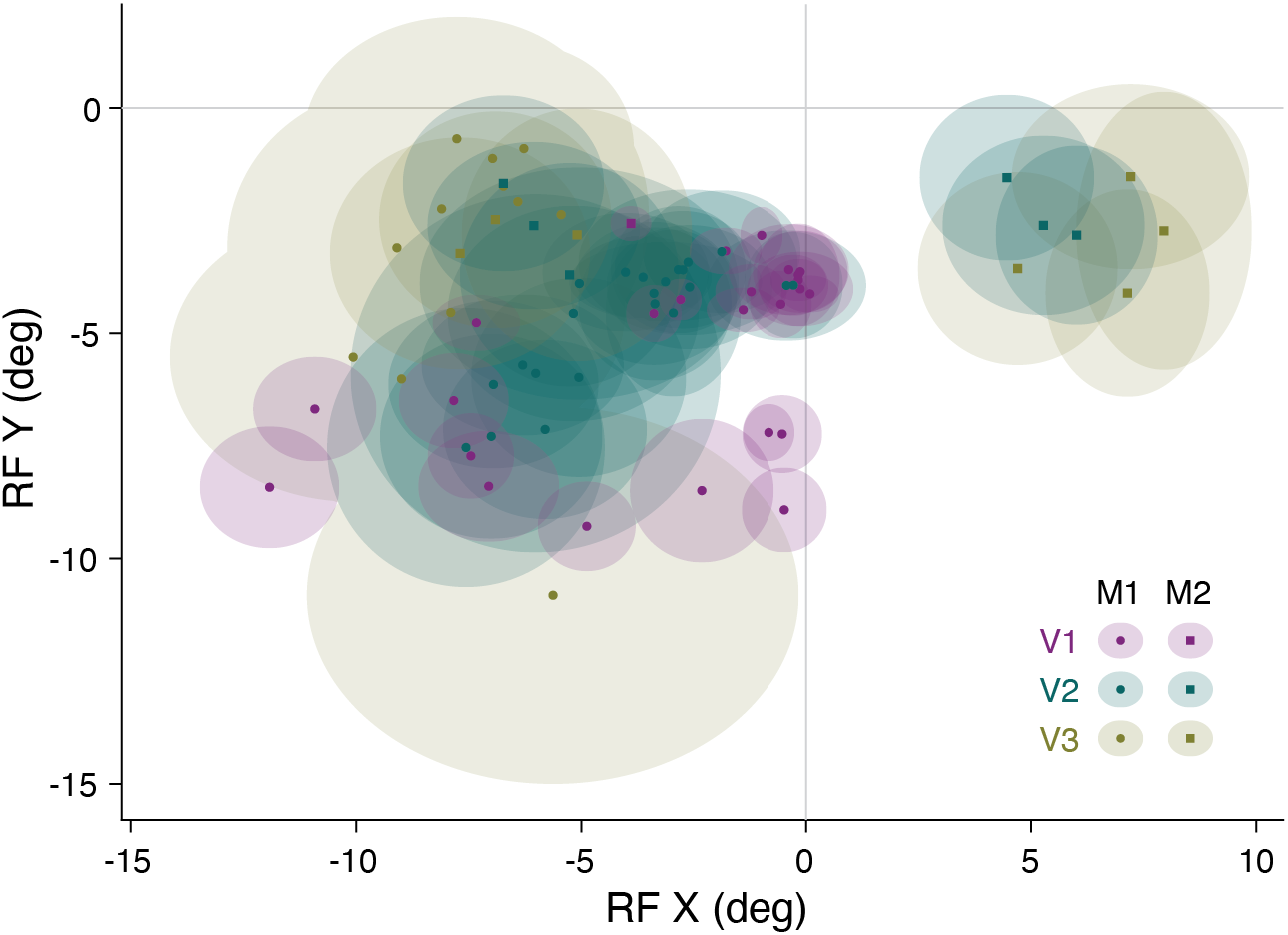
Receptive Field distribution. The average receptive field centers and widths (shaded ellipses) for each session and area are plotted for animal M1 (circles) and animal M2 (squares). The median eccentricity of the receptive fields of the recorded units for V1 was 5.7° (operculum: 4.2° ranging from 2.9° to 7.3°; calcarine sulcus: 10.3°, ranging from 8.2° to 15.1°), 6.2° (range: 3.4° to 11.5°) for V2 and 7.9° (range: 3.1° to 14.0°) for V3/V3A.

**Fig. S2.**
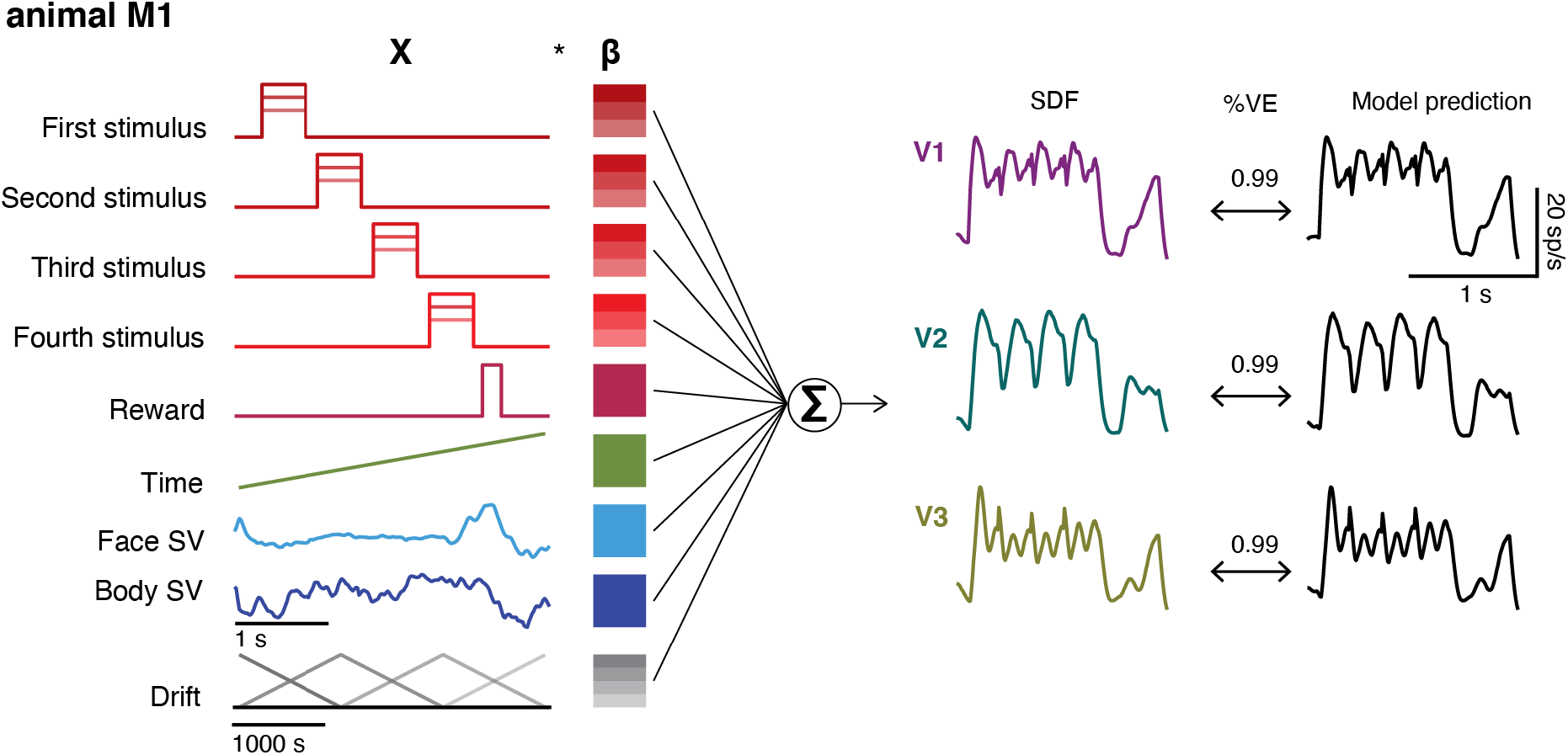
Schematic of the linear encoding model for animal M1. Linear encoding model predicts neural firing in visual cortex (the predictors, labels left, are for the task used in M1). The three traces show peristimulus spike-density function for example units in V1, V2, and V3/V3A (left) recorded in M1, the model predictions (right), and the variance explained by these predictions (center).

**Fig. S3.**
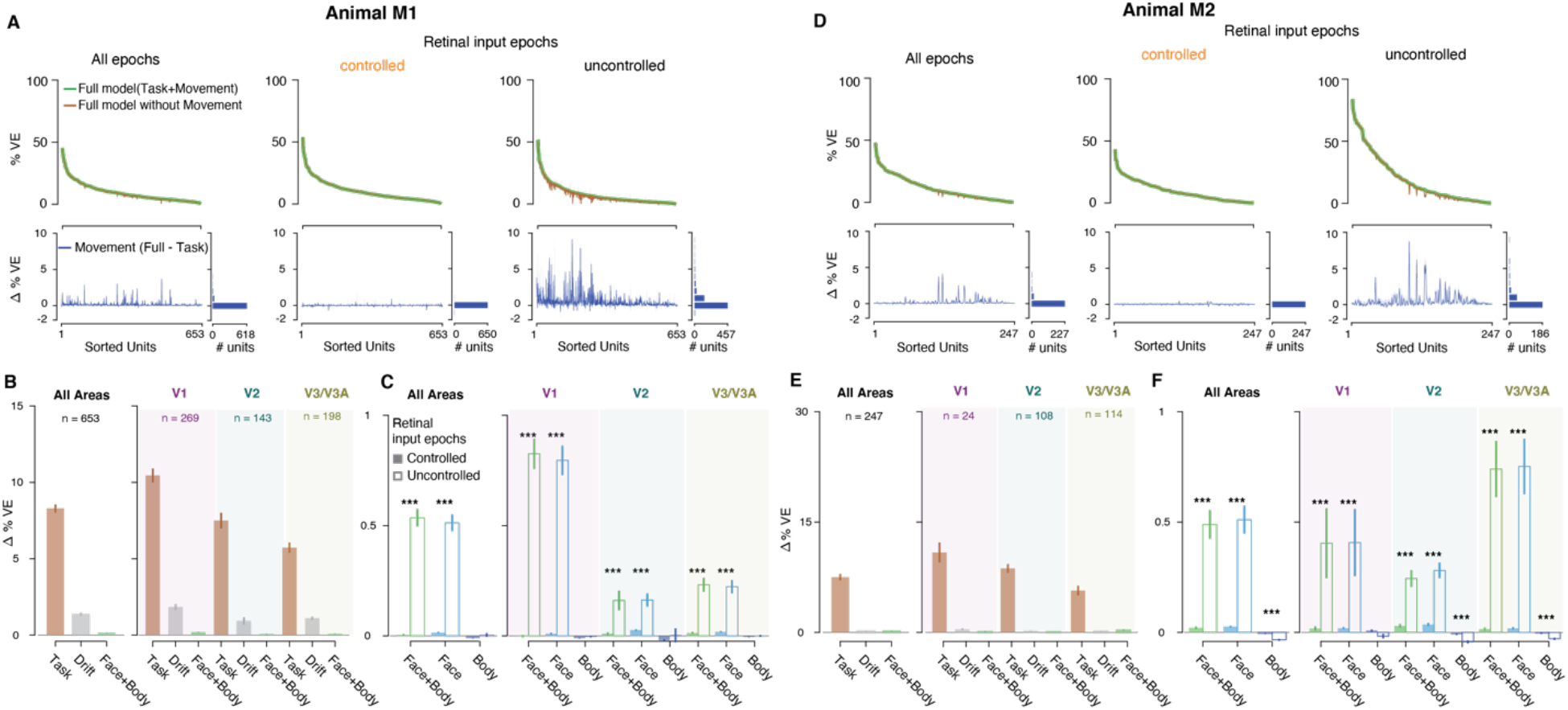
Linear encoding model fits, separately for animals M1(A, B, C) and M2 (D, E, F). **(A, D)** Top: Variance explained by the model with (green), and without (brown) movement covariates for all epochs (left), and separately for epochs when retinal input was controlled (middle), and not controlled (right) for all units. Bottom: Difference in variance explained by the two models. Units are sorted according to the variance explained by the full model. **(B, E)** Unique variance explained by different covariates towards the full model, for units across all areas (left), and separated by area (right). **(C, F)** Unique variance explained by covariates. Format as in **Fig. 2**. For M1: during controlled retinal input epochs (**A**, middle), 3% of units (V1: 15/269, V2: 10/143, V3/V3A: 6/198), and during uncontrolled retinal input epochs, (**A**, right), 55% of units cross the threshold of Δ%VE > 0.1 (V1: 232/269, V2: 67/143, V3/V3A: 108/198); for M2: during controlled retinal input epochs (**D**, middle), 6% of units (V1: 0/24, V2: 6/108, V3/V3A: 7/114), and during uncontrolled retinal input epochs, (**D**, right), 66% of units cross the threshold (V1: 14/24, V2: 64/108, V3/V3A: 83/114).

**Fig. S4.**
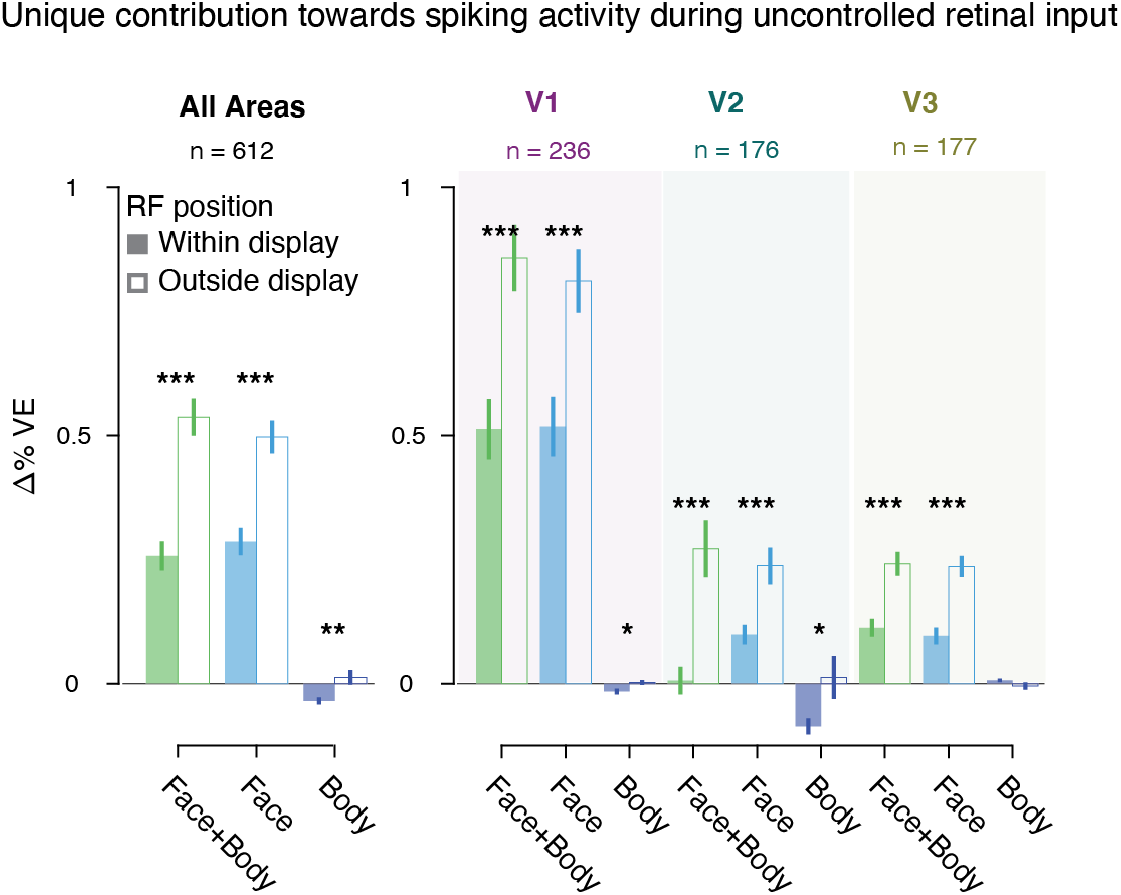
Movements have minimal effect on neural activity after controlling for eye movements in epochs when the animals do not maintain visual fixation. Unique variance explained by different covariates towards the full model during uncontrolled retinal input (open bars in **Fig. 2G**). Unique variance was computed separately for time-points when the receptive field (RF) of the unit was on the monitor showing a gray screen (shaded bars; time-points when retinal input could be inferred), and when the receptive field was outside the boundaries of the monitor (open bars; time-points when retinal input could not be inferred). The criterion for defining whether the RF was on the monitor was that the center of the RF + twice its width was within the monitor edges along the horizontal and vertical dimension. In addition to our general inclusion criteria (see Methods) we required that for each unit the ratio of the number of time-points for which the retinal input could be inferred vs the number of time-points when it could not be inferred, and vice-versa, was at least 10%. This was done to ensure that there were enough time-points for computing unique variance, but our conclusions do not depend on incorporating this additional criterion. Format as in **Fig. 2E**.

**Fig. S5.**
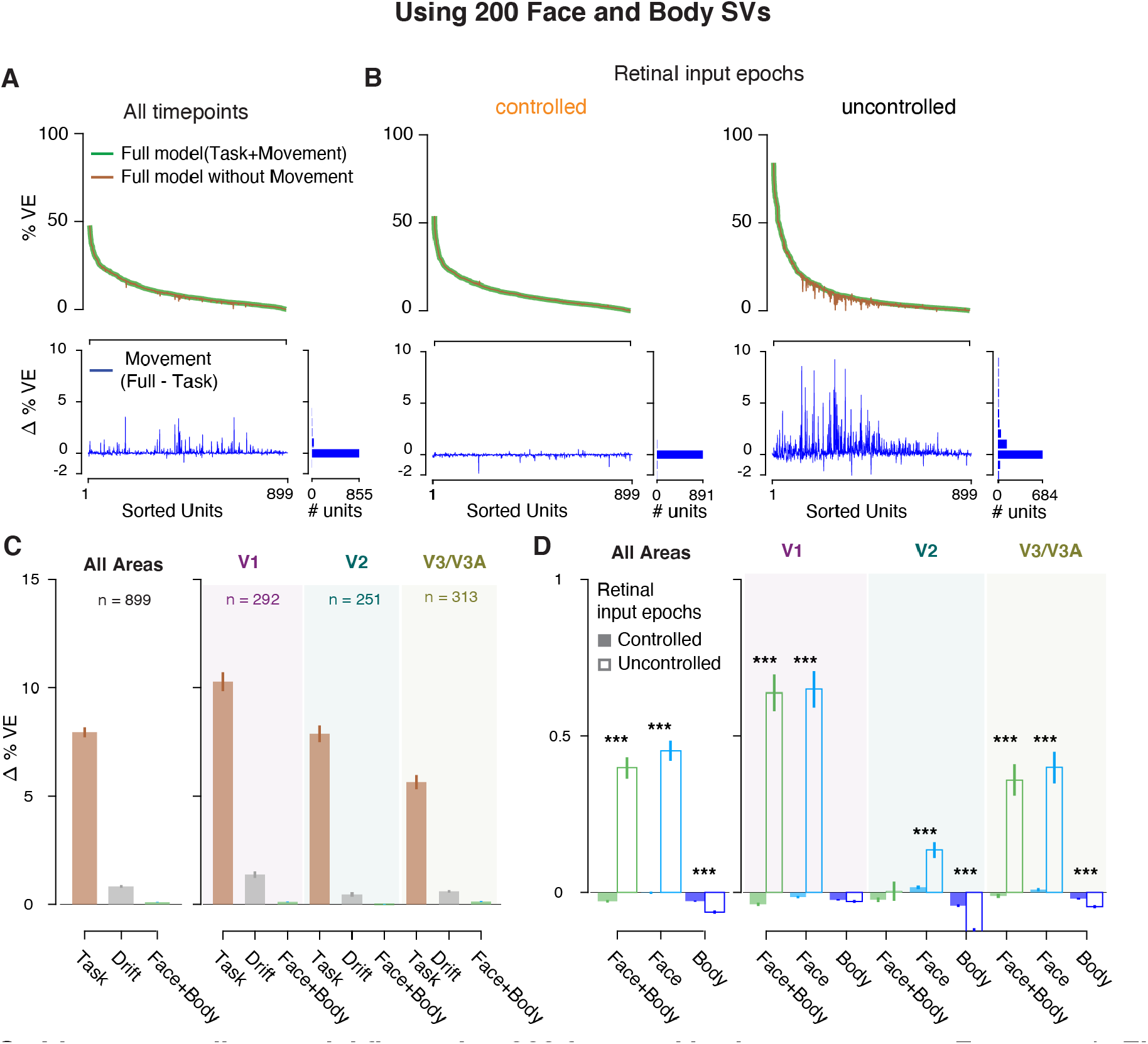
Linear encoding model fits, using 200 face and body components. Format as in **Fig. 2**. Increasing the number of SV components from face and body videos in the linear encoding model to 200 from 30 (**Fig. 2**) did not increase the variance explained by spontaneous movements.

**Fig. S6.**
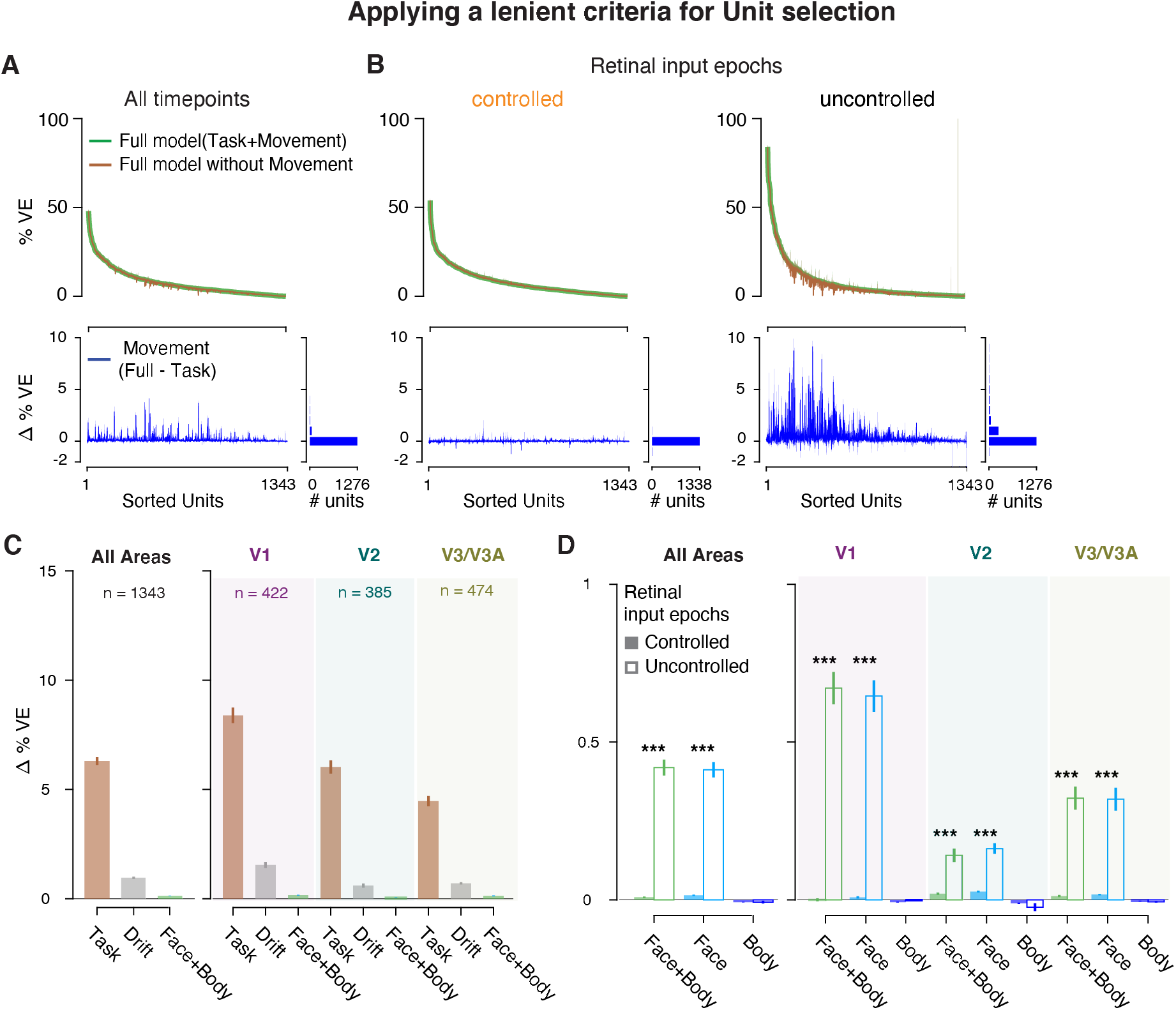
Linear encoding model fits, using a lenient criterion for unit selection. Format as in **Fig. 2**. We used all units for which the % VE by the full model was > 0 (see Methods).

##### Movie S1

Example video clip showing typical spontaneous movements of one of the animals (M1) during a recording session. The labels of the body parts (obtained using DeepLabCut^60^) are only included for demonstration purposes here but not used for our analysis.

**Table S1.** Proportion of units for which the unique variance of movements (Fig **2E**) exceeds different thresholds of unique variance. P-values compare the proportions between controlled and uncontrolled retinal input epochs using a chi-square test.

| Threshold (% ΔVE) | Controlled Retinal input | Uncontrolled retinal input | P-value ($\chi^2$ test) |
| --- | --- | --- | --- |
| All areas |  |  |  |
| 0 | 549/900; 61% | 768/900; 85% | $< 10^{-16}$ |
| 0.05 | 172/900; 19% | 697/900; 77% | $< 10^{-16}$ |
| 0.1 | 48/900; 5% | 606/900; 67% | $< 10^{-16}$ |
| 0.5 | 1/900; 0.1% | 252/900; 28% | $< 10^{-16}$ |
| 1 | 0/900; 0% | 125/900; 14% | $< 10^{-16}$ |
| V1 |  |  |  |
| 0 | 159/293; 54% | 281/293; 96% | $< 10^{-16}$ |
| 0.05 | 42/293; 14% | 268/293; 91% | $< 10^{-16}$ |
| 0.1 | 15/293; 5% | 246/293; 84% | $< 10^{-16}$ |
| 0.5 | 1/293; 0.3% | 132/293; 45% | $< 10^{-16}$ |
| 1 | 0/293; 0% | 67/293; 23% | $< 10^{-16}$ |
| V2 |  |  |  |
| 0 | 160/251; 64% | 168/251; 67% | 0.45 |
| 0.05 | 59/251; 24% | 145/251; 58% | $10^{-15}$ |
| 0.1 | 16/251; 6% | 131/251; 52% | $< 10^{-16}$ |
| 0.5 | 0/251; 0% | 35/251; 23% | $10^{-9}$ |
| 1 | 0/251; 0% | 15/251; 6% | $10^{-4}$ |
| V3/V3A |  |  |  |
| 0 | 211/312; 68% | 277/312; 89% | $10^{-10}$ |
| 0.05 | 64/312; 20% | 242/312; 78% | $< 10^{-16}$ |
| 0.1 | 13/312; 4% | 191/312; 61% | $< 10^{-16}$ |
| 0.5 | 0/312; 0% | 61/312; 20% | $10^{-16}$ |
| 1 | 0/312; 0% | 28/312; 9% | $10^{-9}$ |

## Notes

### Competing Interest Statement

The authors have declared no competing interest.

